# Neurovirulent and non-neurovirulent strains of HIV-1 and their Tat proteins induce differential cytokine-chemokine profiles

**DOI:** 10.1101/2024.11.03.621750

**Authors:** Vasudev R. Rao, Milka Rodriguez, Siddappa N. Byrareddy, Udaykumar Ranga, Vinayaka R. Prasad

**Affiliations:** Department of Microbiology and Immunology, Albert Einstein College of Medicine, Bronx, NY, USA; Department of Pharmacology and Experimental Neuroscience, University of Nebraska Medical Center, Omaha, NE, USA; Molecular Biology & Genetics Unit, Jawaharlal Nehru Centre for Advanced Scientific Research, Bangalore, India

**Keywords:** HIV neuropathogenesis, Immune response, viral pathogenesis, inflammation, Tat, Inflammatory cytokines

## Abstract

HIV-1 enters the central nervous system (CNS) early in infection, and a significant proportion of people with HIV experience CNS complications despite anti-retroviral therapy. Chronic immune dysfunction, inflammatory cytokines and chemokines, and viral proteins like Tat and gp120 released by HIV-1-infected immune cells are implicated in the pathogenesis of HIV-1-associated neurocognitive disorders (HAND). To elucidate the contribution of non-viral factors to CNS complications in people with HIV-1, a comparative analysis of neurovirulent subtype B (HIV-1_ADA_) and non-neurovirulent subtype C (HIV-1_Indie-C1_) isolates was performed. Culture supernatants from HIV-1-infected PBMCs, either with or without immunodepletion of Tat and gp120, were used to treat SH-SY5Y neuroblastoma cells. HIV-1_ADA_-infected PBMC media showed significantly higher cytotoxicity than HIV-1_IndieC1_-infected _PBMC_ media, notwithstanding the depletion of Tat and gp120, highlighting the role of non-viral factors contributing to neurotoxicity. A comparison of inflammatory profiles revealed that HIV-1_ADA_ media contained elevated levels of cytokines (IL-1α, IL-1β, IL-6, TNFα) and chemokines (CCL2, CCL3, CCL4, IP10). Given the involvement of Tat in upregulating immune mediators, PBMCs from healthy subjects were treated with recombinant purified Tat from subtype B or C. Subtype B Tat induced a stronger inflammatory response than subtype C Tat. These results confirm that both viral and non-viral immune factors mediate neuronal damage in people with HIV.

## Introduction

HIV-associated neurocognitive disorders (HAND) comprise a spectrum of neurodegenerative pathologies that arise from the viral infection of the central nervous system (CNS) in subjects with chronic HIV infection. Despite the success of combination of antiretroviral therapy (cART), HAND persists as a significant clinical challenge in up to 50% of HIV-infected individuals, with its adverse effects on patient survival, quality-of-life metrics, and activities of daily living (1). While subjects on cART show low levels of systemic and CNS viral replication, the residual viral proliferation can induce the secretion of several pro-inflammatory cytokines and chemokines that, in turn, cause augmented neurological damage. Several chemokines and cytokines, including IP-10, IL-1α, IL-1β, and TNF-α, have been implicated in the etiology of HAND and detected at elevated levels in CSF and serum in patients suffering from HAND (2). Although most studies primarily focused on macrophages and microglia as the source of secreted chemokines and cytokines, other reports demonstrated the migration of both HIV-1-infected T-cells and monocytes to the CNS, implicating their role in neuronal damage in HAND (3, 4).

This prompted us to employ peripheral blood mononuclear cells (PBMCs), consisting of both macrophages and T-cells, that were infected with relatively low titers of HIV-1 and examine the contribution of non-viral neurotoxic factors to HAND. Neurotoxic factors other than those of viral origin (e.g., Tat and gp120), are present in detectible but low levels in the CNS of people with HIV (PWH) who are virologically suppressed *via* cART. Previously, we demonstrated that intra- and inter-subtype genetic variations in HIV-1 Tat and gp120 influence neurovirulence (5-7) we evaluated the effects on neurocognitive functions in a SCID-HIV encephalitis mouse model. Using cell models, we monitored neurotoxicity, pro-inflammatory cytokine induction, and monocyte recruitment. Our previous comparative analyses demonstrated significantly higher levels of neurovirulence by HIV-1_ADA_ (a representative neurovirulent strain of subtype B) compared to HIV-1_IndieC1_ (a representative non-neurovirulent strain of subtype C) (7, 8).

In this study, using HIV-infected PBMCs, we compare the relative effects of chronic inflammation and immune dysfunction mediated by HIV-1_ADA_ and HIV-1_Indie_ infections to HAND. Using spent media of PBMC infected with either of the viral strains, we ascertain the augmented neurotoxic properties of HIV-1_ADA_ over HIV-1_Indie_. Additionally, after immunodepleting Tat and gp120 proteins from the spent media, we show sustained, although reduced, neurotoxic properties of the HIV-1_ADA_, which we ascribe to pro-inflammatory cytokines and chemokines secreted into the media. Furthermore, we demonstrate that a higher level of induced secretion of pro-inflammatory cytokines and chemokines by treating PBMC with recombinant Tat proteins derived from HIV-1 subtypes B when compared to that from subtype C. Thus, from our data, that pre-deterministic role of Tat in promoting neurotoxicity is evident and this property appears to be subtype-specific.

## Materials and Methods

### Materials

PBMCs were purified from leukopaks from New York Blood Center. Human monocyte-derived macrophages (MDMs) were obtained by differentiating CD14^+^ monocytes isolated from peripheral blood mononuclear cells (PBMCs). Recombinant purified Tat proteins from HIV-1_BL43_ (9), a subtype C virus and HIV-1_YU2_, a subtype B virus were generated as described previously (9).

### Isolation of peripheral blood mononuclear cells

Peripheral blood mononuclear cells (PBMCs) were isolated from HIV-1 negative donor leukopaks (New York Blood Center) using density gradient centrifugation. Leukopaks were diluted in RPMI/FBS and overlaid over a Ficoll-Hypaque (Sigma) gradient. Following centrifugation at 400 xg for 30-40 min at room temperature with no brakes, PBMCs were recovered from the Ficoll-Hypaque interface, diluted in PBS, and centrifuged at 200 xg for 10 min at room temperature to eliminate platelets. PBMCs were re-suspended in RPMI (Gibco), supplemented with 10% FBS (Gibco) and 1% Penicillin and Streptomycin. Cells were incubated at 37°C in T75 flasks.

### HIV-1 infection of PBMCs

PBMCs were activated with PHA (5 µg/ml) and IL-2 (100 IU/ml) for 48 hours prior to infection. HIV-1_ADA_ and HIV-1_IndieC1_ virus stocks were generated by transfecting the HEK 293T cells with infectious molecular clone DNAs. Virus released into the medium was quantitated by the p24 Enzyme-linked immune-sorbent assay (ELISA; Applied Biosystems, Inc), as described previously (6). Approximately 50 x 10^6^ PBMCs were infected with viral stocks at low multiplicity of infection (5 ng and 10 ng of HIV-1_ADA_ and HIV-1_IndieC1_, respectively) to achieve comparable levels of viral infectivity at the assay end point 4-hours or five days following infection as described previously (6).

### Immunodepleting Tat and gp120

HIV-1 Tat and gp120 were immuno-depleted from HIV-infected spent media from PBMCs using broadly neutralizing anti-gp120 (VRC 03, NIH AIDS Repository) and anti-Tat E1.1 (9) antibodies, as described previously (6, 7). For this purpose, Pansorbin beads (Calbiochem, Cat no. 507861) were incubated with anti-gp120 and anti-Tat antibodies for an hour on ice. Antibodies were used in excess, at 3 -100 fold higher concentrations than required (VRC03 and E1.1 antibodies were used at 150 to 300 ng and 300 ng, respectively) to ensure complete protein depletion from spent media. The beads were centrifuged, re-suspended in 25 μL RPMI, and added to 10 ml of spent media. Following incubation for an hour on ice, the Pansorbin beads are centrifuged at 3,000 rpm for 5 minutes at 4°C. The antigen-depletion step was repeated to ensure the complete removal of gp120 and Tat proteins from the supernatant. The antibody-treated supernatants were used in the *in vitro* neurotoxicity assays.

### Neurotoxicity assay

SH-SY5Y neuroblastoma cells were obtained from ATCC (CRL-2266) and maintained in a 1:1 mixture of Eagle’s Minimal Essential Medium and F12 Medium supplemented with 10% fetal bovine serum. Approximately 250,000 cells/well were plated in 12-well plates and allowed to stabilize for three days. Three hundred µl (300 µL) of HIV-infected PBMC supernatants (with or without immuno-depletion) were added to 600 µL of SH-SY5Y media and then added to previously plated SH-SY5Y cells for 24 hours as described previously (6). WST-1 assay (Roche) was used to determine neuronal viability in triplicate wells. The percentage of cell death, compared to control cells treated with uninfected PBMC supernatant, was calculated as previously described (6).

### Luminex ELISA and CCL2 ELISA assays

Thirteen different cytokines/chemokines (Interferon-α, CCL-2, CCL-3, CCL-4, IL1-α, IL-1β, IL-4, IL-6, IL-8, IL-10, TNF-α, IP-10 and RANTES) were measured using a high-sensitivity human cytokine array kit. Each sample was assayed in triplicate wells, and the cytokine standards supplied by the manufacturer were included on each plate. Data was acquired using a Luminex-100 system and analyzed using the Bio-Plex Manager software. As CCL2 levels were above the limit of detection of the cytokine array kit, a separate CCL-2 ELISA was performed using the Invitrogen Human CCL2 ELISA kit, as described previously (7, 8).

### Tat treatment of PBMCs and Real-time PCR

Purification of recombinant HIV-1 Tat proteins from the neurovirulent Subtype-B YU-2 strain and non-neurovirulent Subtype-C BL-43 strain has been reported previously (9). Neurotoxic properties of these Tat proteins were described previously (10). PBMCs of healthy subjects were activated with PHA (5 µg/ml) for 24 hours, before treating them with 200ng/ml recombinant Tat protein for 4 hours at 37°C. PBMCs were pelleted, and RNA was extracted using RNAeasy Kit (Qiagen). cDNA was generated from the RNA using a Taqman reverse transcriptase cDNA kit (Applied Biosystems Inc). cDNA was quantitated, and a qRT-PCR was performed using a Taqman cytokine gene expression array plate (Applied Biosystems Inc). The Delta delta-CT method was used to calculate fold differences between treatments, using GAPDH as a control house-keeping gene. RNA extracted from PBMCs treated with a buffer was used as a control, and the relative fold changes were determined in relation to control CT values.

## Results

### Contribution of non-viral factors to HIV neuropathogenesis *in vitro*

To understand the contribution of non-viral immune factors to the pathogenesis underlying HAND, we investigated whether HIV-spent media immunodepleted for viral Tat and gp120 proteins could still retain neurotoxicity. PBMCs of healthy donors were infected with the neurovirulent HIV-1_ADA_ (5 ng/ml p24) or the non-neurovirulent HIV-1_Indie_ (10 ng/ml p24) strain to achieve comparable levels of viral infection five days after infection. Following viral infection, p24 levels in the spent media were quantitated using a commercial p24 assay. Spent media of HIV-1_ADA_ and HIV-1_Indie_ contained 25ng/ml and 22ng/ml of p24, respectively, confirming comparable levels of infection. Spent media were then incubated with VRC03 and E1.1 antibodies to deplete gp120 and Tat antigens from the samples, respectively. Subsequently, SHSY-5Y neuroblastoma cells were treated with immuno-depleted supernatants or an uninfected control medium. Following immuno-depletion of Tat and gp120 (the two primary viral factors responsible for neurotoxicity), the neurovirulent HIV-1_ADA_ isolate retained about half of its neurotoxicity compared to the non-neurovirulent HIV-1_IndieC1_ isolate (Figure 1), that exhibited barely any change in its already low overall neurotoxicity (Figure 1 inset). This result pointed at host factors other than viral proteins, such as pro-inflammatory cytokines and chemokines, as the potential source of cytotoxicity of SHSY-5Y cells.

**Figure 1:**
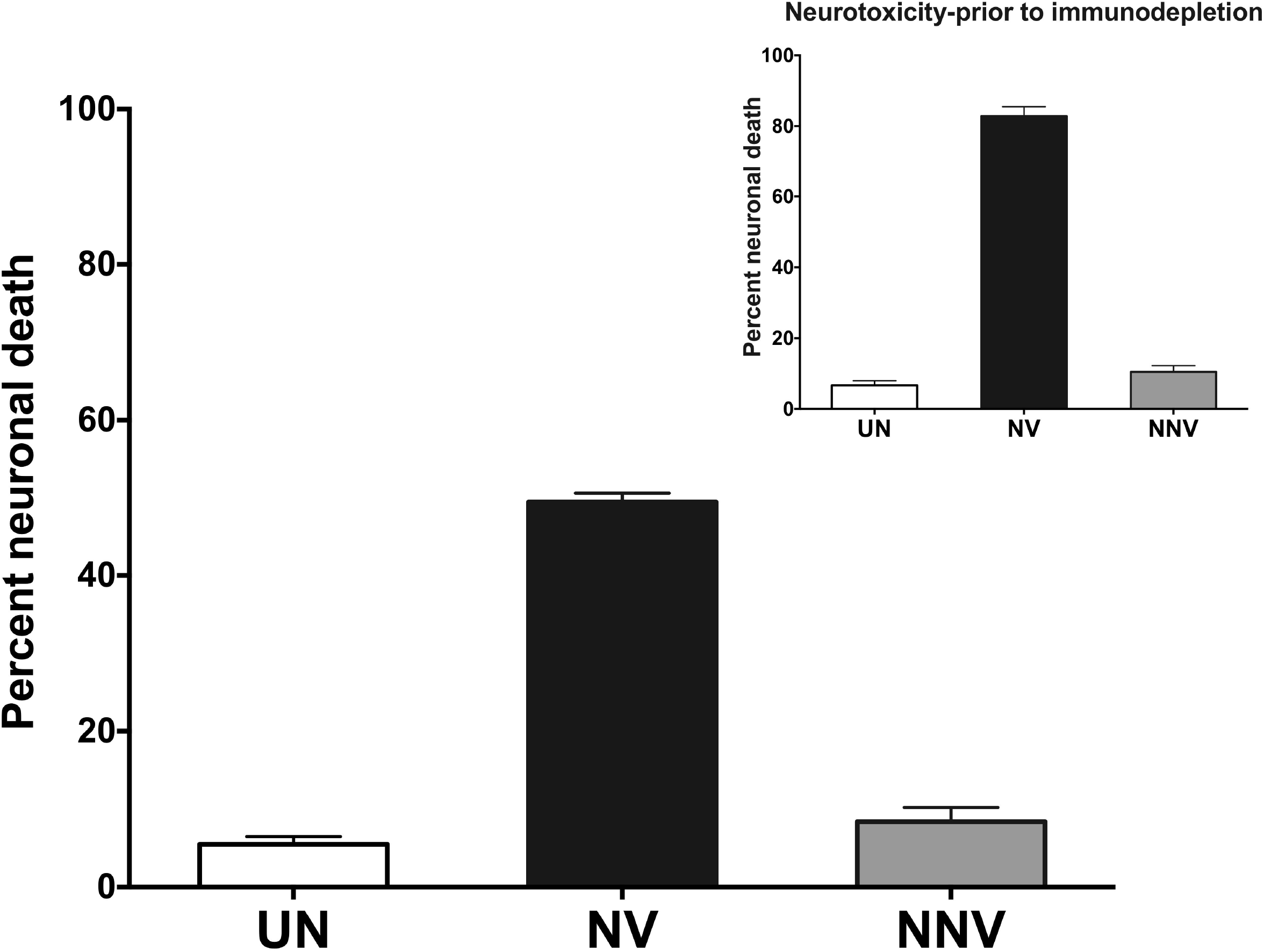
Contribution of cytokines and chemokines to neurotoxicity: 300 µl of the spent media of PBMC infected with HIV-1_ADA_ or HIV-1_Indie_, were incubated with pooled VRC03 and E1.1 (anti-gp120 and anti-Tat, respectively) antibodies for 24 h to deplete both gp120 and Tat, via two rounds of immunodepletion. Neuronal SH-SY5Y cells were incubated with the spent media, with or without immuno-depletion. WST-1 cell viability was performed using the WST-1 assay (Inset-adapted from reference #6).

### Differential induction of cytokines and chemokine by HIV-1_ADA_ and HIV-1_IndieC1_

Upon infection of PBMCs, HIV-1_ADA_ induced significantly higher levels of pro-inflammatory cytokines IL-1α, IL-1β, IL-6 and TNF-α (Figure 2). These factors have been implicated (6, 11-15) in causing neuronal damage in various neuroimmune disorders. In contrast, HIV-1_IndieC1_ induced a higher level expression of IL-8, which is an exception in not being a neurotoxic cytokine. Further, HIV-1_ADA_ also induced higher levels of CC family of chemokines, CCL2, CCL3, and CCL4, and also IP-10 (CXCL-10), a CXC family chemokine. Although the CC-chemokines are conditionally neuroprotective (16, 17), at higher levels, they can promote increased immune cell trafficking to the brain, leading to augmented neuronal damage. Furthermore, HIV-1_ADA_ infection of PBMCs also resulted in a more significant down-regulation of anti-inflammatory cytokines IL-4 and IL-10, compared to HIV-1_IndieC1_-infection or control uninfected sample. Additionally, HIV-1_IndieC1_ also suppressed the induction of IL-10 to a greater extent than that of HIV-1_ADA_. These results suggest that the cytokines and chemokines are vital in causing neurotoxicity, which is subjective to subtype-specific differences.

**Figure 2:**
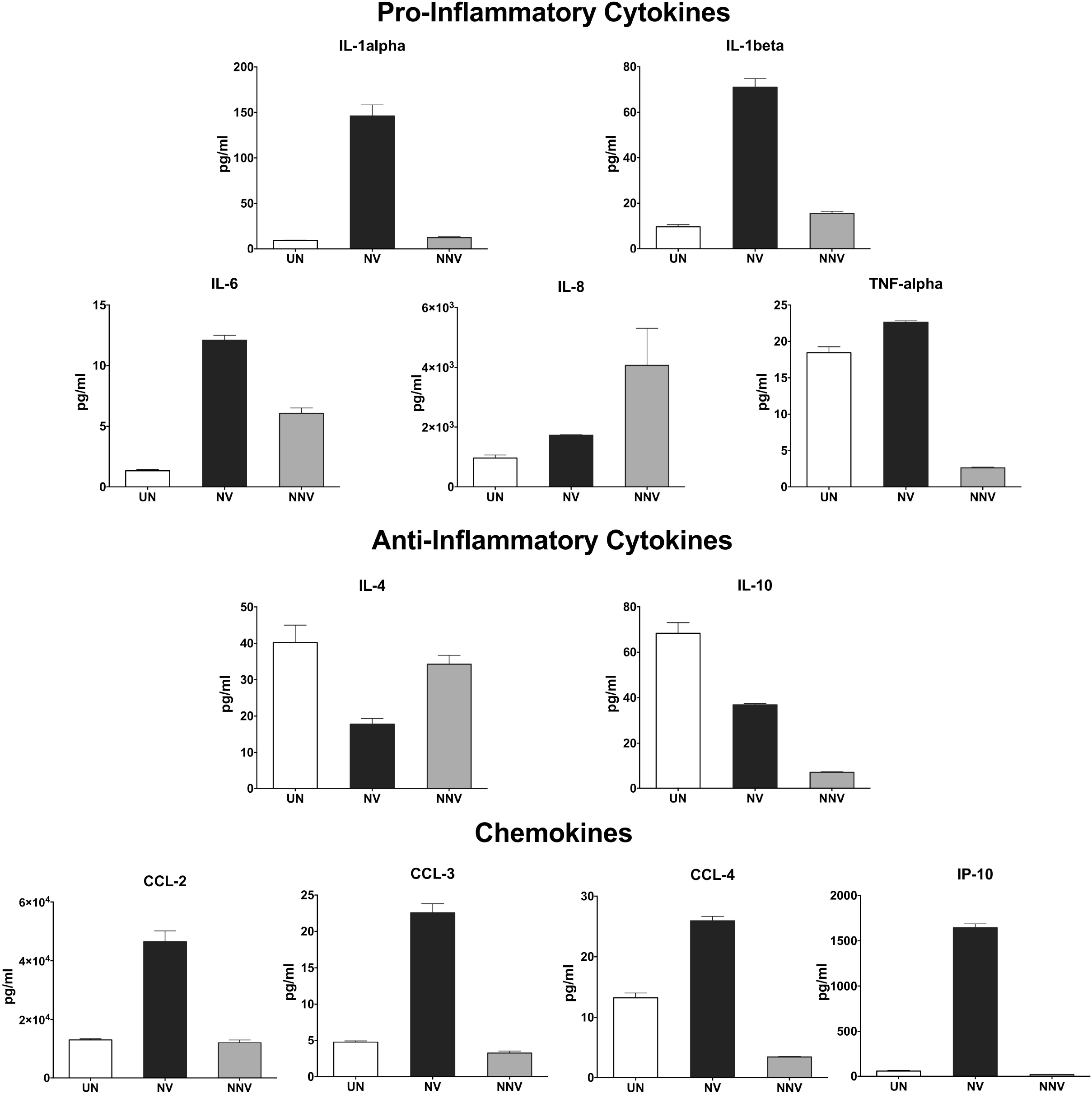
Differential induction of chemokines and cytokines upon HIV-1 infection of PBMCs. A panel of thirteen cytokines/chemokines (Interferon-α, CCL-2, CCL-3, CCL-4, IL-1α, IL-1β, IL-4, IL-6, IL-8, IL-10, TNF-α, IP-10, and RANTES) was measured in the spent media from HIV-1_ADA_- or HIV-1_Indie_ -infected PBMCs, using a high-sensitivity human cytokine array kit. Each sample was assayed in triplicate wells, and data was acquired using a Luminex-100 system and analyzed using the Bio-Plex Manager software.

### Tat plays a key role in the differential induction of chemokines and cytokines

To further characterize the neurotoxic effects of Tat, known for modulating the expression of cellular inflammatory genes, PBMCs were treated with recombinant Tat proteins from different strains HIV-1_YU-2_ and HIV-1_BL-43_, representing subtypes B and C, respectively. After a 4-hour treatment, RNA was extracted from PBMCs, and gene expression changes were compared using quantitative real-time PCR on a macroarray of selected cytokine/chemokine genes. The neurovirulent HIV-1_YU-2_ Tat protein induced 5-30 fold increases in the level of pro-inflammatory cytokine transcripts compared to the control or the non-neurovirulent HIV-1_BL-43_ Tat (Figure 3). Similar elevated patterns for subtype B Tat-treated PBMCs were observed with neurotoxic host factors, such as TNF-α and IP-10.

**Figure 3:**
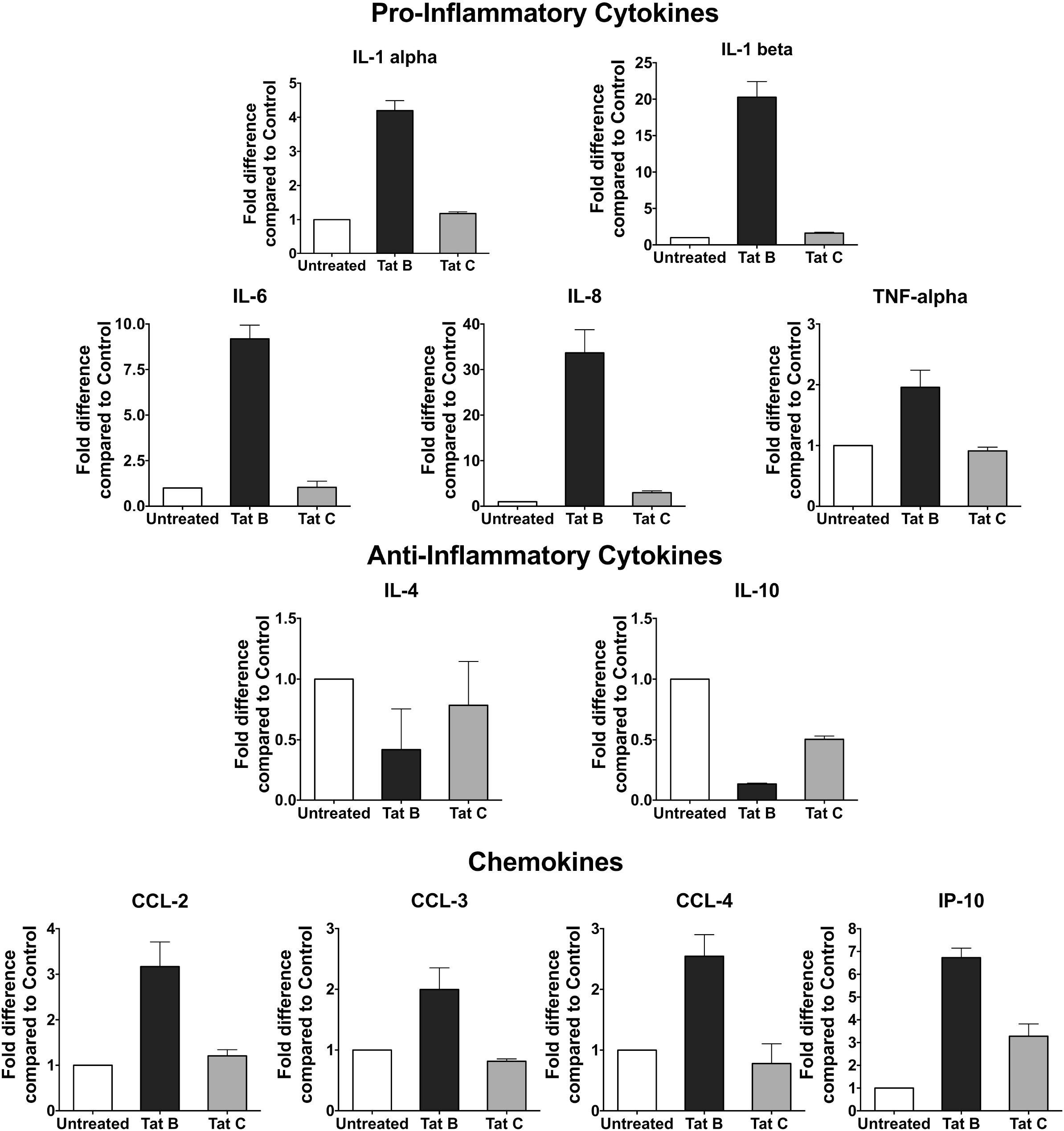
Differential secretion of cytokines and chemokines by PBMC treated with Tat: PBMCs of healthy donors were treated with 200 ng/ml of recombinant Tat proteins of HIV-1_YU-2_ (clade B) or HIV-1_BL-43_ (clade C) viruses. Transcription of selected cytokine and chemokine genes was evaluated using real-time PCR.

The Tat-induced patterns of the anti-inflammatory cytokines, IL-4 and IL-10, differed from those of the pro-inflammatory cytokines. Both Tat proteins exerted a suppressive effect on the expression of the two anti-inflammatory cytokines, compared to a ‘no Tat’ control. Importantly, HIV-1_BL-43_ Tat exerted a weaker inhibitory effect on the secretion of these two anti-inflammatory cytokines than the HIV-1_YU-2_ Tat, thus inducing still significantly higher levels of these host factors. Thus, compared to subtype B, subtype C Tat induced relatively low levels of pro-inflammatory cytokines and relatively higher levels of anti-inflammatory cytokines, resulting in an overall lower potential for neurotoxicity

## Discussion

Neurodegeneration persists in HIV-infected individuals despite cART and viral suppression due to multifactorial etiopathogenesis (Winston and Spudich, 2020). Neurotoxic cytokines and chemokines secreted by HIV-1 infected cells and bystander cells play a central role in developing neuronal damage observed in HAND (2). Using *in vitro* HIV-infection of macrophages and the measurement of neurocognitive functions in SCID-HIVE mice, we previously established differential neurovirulent properties of HIV-1 B and C subtypes and furthermore demonstrated that subtype-specific genetic variations in Tat or gp120 primarily underlie the observed differences in neurotoxicity.

### A potential role of non-viral factors in neuroinflammation

The spent media of virus-infected PBMC, retained neurotoxic properties even after depleting Tat and gp120, suggesting an important role for immune dysregulation in the causation of adverse CNS outcomes. Using the highly sensitive Luminex assay, several pro- and anti-inflammatory cytokines and chemokines could be measured in such media. Importantly, the neurovirulent subtype B strain up-regulated the cytokines higher than the non-neurovirulent subtype C strain. Consistent with these results, differential expression of the same cytokines was also shown in PBMCs treated with recombinant Tat proteins.

### Both purified Tat proteins and HIV-infection display subtype-specific differences

Our observations with HIV-infected PBMCs correlated well with the findings using recombinant purified Tat proteins added to PBMCs. Subtype C Tat protein induced low levels of pro-inflammatory cytokines than subtype B Tat protein and higher levels of anti-inflammatory cytokines. Furthermore, previously published studies (17) (18, 19) on IL-10 levels induced by the subtype B and C Tat proteins in monocytes have yielded inconsistent results suggesting the involvement of additional viral or host factors affecting IL-10 regulation. It is well known that Tat protein is secreted and is taken up by bystander cells (2). In the uninfected bystander cells, HIV-1 Tat can transcriptionally transactivate proinflammatory cytokine genes. We previously showed that the differential uptake of Tat B and C proteins is likely the basis of differential transactivation of such inflammatory host genes and the resulting differential neuroinflammation (20). These observations indicate that differences between neurovirulent and non-neurovirulent HIV-1 strains are reflected in their Tat protein, inducing the differential regulation of pro- and anti-inflammatory chemokines/cytokines in PBMCs.

Among the factors up-regulated, both IL-1α and IL-1β have been implicated in causing and enhancing neuronal damage (21). These variant forms of Interleukin-1 act through the IL-1 receptor. Several models of brain injury demonstrated that IL-1 receptor agonist (IL-1Rα) reduces neuronal damage caused by IL-1α and IL-1β (11). IL-6 has been implicated in enhancing NMDA-mediated neurotoxicity, leading to increased membrane depolarization (22). In the CNS, TNF-α orchestrates a diverse array of functions, including but not limited to immune surveillance, cellular homeostasis, and protection against specific neurological insults. However, in the context of HIV-1 infection, TNF-α enhances HIV-1 Tat-induced neurotoxicity by oxidative stress leading to neuronal apoptosis (23). TNF-α can also inhibit glutamate uptake by astrocytes, leading to an extracellular buildup of glutamate, causing neuronal excitotoxicity (24). Furthermore, inhibiting TNF-α activity abrogates HIV-1-induced neurotoxicity *in vitro* (25). IL-1β and TNF-α dysregulate glutamate production in neurons through the induction of glutaminase (15). IP-10, while serving as a T cell chemoattractant, can cause neurotoxicity in primary neurons due to elevated intracellular calcium levels (12), leading to caspase activation and neuronal death (26).

We found that infection of PBMC by the neurovirulent HIV-1_ADA_ and treatment of PBMC by recombinant Tat of HIV-1_YU-2_ strain, both up-regulated chemokines CCL-2, CCL-3, and CCL-4 to significant levels. CCL-2 (MCP-1), although typically neuroprotective (16), can cause monocyte migration to the CNS (27), thus exacerbating neuronal damage. Likewise, CCL-3 (MIP1α) and CCL-4 (MIP1β) also can enhance the migration of immune cells of diverse lineages to the CNS (28). All the processes can contribute to increased chronic CAN inflammation and progressive neuronal damage, finally manifesting as HAND. Neuronal injury is further augmented by the suppression of anti-inflammatory chemokines, IL-4 and and IL-10, as shown *in vitro* in neuronal cultures and *in vivo* in patients. Despite viral suppression with cART, indirect neuronal damage is persistent and is mediated by upregulated inflammatory cytokines and chemokines, particularly in individuals with chronic HIV infection. Concomitant up-regulation of several pro-inflammatory cytokines, such as TNF-α and IL-1α, and down-regulation of anti-inflammatory cytokines, such as IL-4 and IL-10, have been documented in HIV-1 patients suffering from HAND (13).

Our findings suggest that in addition to using cART to control viral replication, adjunctive therapies incorporating neuroprotective agents, such as minocycline (29, 30) and memantine (31) might be beneficial in suppressing CNS inflammation. Alternative approaches to reducing CNS inflammation by targeting leukocyte trafficking across the blood-brain barrier with Natalizumab, an anti-alpha-4-integrin monoclonal antibody (32, 33) can potentially improve CNS outcomes in people with HIV. Additionally, novel therapies targeting HIV-1 Tat (34) could reduce overall neuronal damage, both from the direct neurotoxicity of Tat and the up-regulation of pro-inflammatory cytokines and chemokines induced by Tat. Although substantial in vitro and in vivo evidence demonstrates attenuation of neuronal damage using adjunctive approaches, therapeutic interventions targeting CNS dysfunction in HIV-infected individuals remain clinically ineffective, necessitating further research to develop interventions that can improve patient outcomes and quality of life.

## Acknowledgements

The results reported in this manuscript were supported by NIH grants R01 MH083570 and R01 AI153008 to VRP. MAR wishes to acknowledge support from the institutional training grant NIH T32 AI007501. The authors wish to thank the Einstein-Rockefeller-CUNY Center for AIDS research for use of Biomarkers and Advanced Technologies core services; Arthur Ruiz and David Ajasin for critically reading the manuscript and the NIH AIDS Reagent Program, Division of AIDS, NIAID, NIH for providing valuable reagents.

## Author contributions

VR generated the neurotoxicity data, helped draft the original manuscript and helped develop the overall content, MR generated the data on the infection of PBMCs and measurement of cytokine and chemokine transcript levels, SB purified the Tat proteins and generated the data on the cytokine protein levels from PBMCs treated with Tat, UR supervised the work on Tat purification and cytokine protein quantitation and for the manuscript revisions, VR helped develop the overall content, supervised most of the work reported here and contributed to drafting the original manuscript. All authors have accepted responsibility for the entire content of this manuscript and approved its submission.

